# Assessment of immunogenicity and protective efficacy of ZyCoV-D DNA vaccine candidates in Rhesus macaques against SARS-CoV-2 infection

**DOI:** 10.1101/2021.02.02.429480

**Authors:** Pragya D Yadav, Sanjay Kumar, Kshitij Agarwal, Mukul Jain, Dilip R Patil, Kapil Maithal, Basavaraj Mathapati, Suresh Giri, Sreelekshmy Mohandas, Anita Shete, Gajanan Sapkal, Deepak Y Patil, Ayan Dey, Harish Chandra, Gururaj Deshpande, Nivedita Gupta, Dimpal Nyayanit, Himanshu Kaushal, Rima Sahay, Anuradha Tripathy, Rajlaxmi Jain, Abhimanyu Kumar, Prasad Sarkale, Shreekant Baradkar, Chozhavel Rajanathan, Hari Prasad Raju, Satish Patel, Niraj Shah, Pankaj Dwivedi, Dharmendra Singh, Priya Abraham

## Abstract

Vaccines remain the key protective measure to achieve herd immunity to control the disease burden and stop COVID-19 pandemic. We have developed and assessed the immunogenicity and protective efficacy of two formulations (1mg and 2mg) of ZyCoV-D (a plasmid DNA based vaccine candidates) administered through Needle Free Injection System (NFIS) and syringe-needle (intradermal) in rhesus macaques with three dose vaccine regimens. The vaccine candidate 2mg dose administered using Needle Free Injection System (NFIS) elicited a significant immune response with development of SARS-CoV-2 S1 spike region specific IgG and neutralizing antibody (NAb) titers during the immunization phase and significant enhancement in the levels after the virus challenge. In 2 mg NFIS group the IgG and NAb titers were maintained and showed gradual rise during the immunization period (15 weeks) and till 2 weeks after the virus challenge. It also conferred better protection to macaques evident by the viral clearance from nasal swab, throat swab and bronchoalveolar lavage fluid specimens in comparison with macaques from other immunized groups. In contrast, the animals from placebo group developed high levels of viremia and lung disease following the virus challenge. Besides this, the vaccine candidate also induced increase lymphocyte proliferation and cytokines response (IL-6, IL-5).The administration of the vaccine candidate with NFIS generated a better immunogenicity response in comparison to syringe-needle (intradermal route). The study demonstrated immunogenicity and protective efficacy of the vaccine candidate, ZyCoV-D in rhesus macaques.

## INTRODUCTION

The ongoing pandemic of Coronavirus disease-19 (COVID-19) has affected countries across the globe and millions of people have been impacted leading to loss of life, economy and productivity. This has led to development of various intervention strategies including development of vaccine for prophylactic use, monoclonal antibodies and other biologics for therapeutic application.Two mRNA-based vaccines (1) have been granted emergency use authorization (EUA) in UK, USA, Canada, European Union and many other countries across the world. Two adenovirus based non-replicating viral vaccine (Oxford and Sputnik) has been granted EUA by UK, India and Russia. However, the need for additional vaccine candidates still persists to so that vaccines would be available to a larger population across the world. Studies have utilized various vaccine development platforms such as inactivated whole virion, live-attenuated, nucleic acid-based (RNA, DNA), replicating /non-replicating viral vector-based, protein subunit and numerous other platforms (2). The response of the global scientific community has resulted in development of over 233 vaccine candidates until December 29 2020. Of these, 172 candidates are in pre-clinical stage and 61 candidates have progressed to clinical trials (3). One of thesafe and efficacious next generation vaccines is the DNA vaccine which involves direct administration of plasmid DNA encoding for immunogenic antigen component of the pathogen (4). DNA vaccines are composed of bacterial plasmids with a gene encoding for the protein of interest and transcription promoter and terminator. Routes of administration of DNA vaccines can be topical application, parenteral or cytofectin-mediated method. The plasmid gains entry in multiple cells such as myocytes, keratinocytes, and antigen-presenting cells (APCs) where it enters in the nucleus as an episome; without getting integrated into the host cell DNA. Using the host cell’s transcription and translation machinery, the inserted gene gets translatedinto antigen. Protein expressed by plasmid-transfected cells is likely to be expressed within the cell and folded in its native configuration. The antigen is recognized by antigen presenting cells (APCs) and further induces antibodies and cellular response. DNA vaccines are free of any infectious agent, are low cost, fairly safe and relatively easy for large-scale manufacturing (5). They induce strong and long-lasting humoral and cell-mediated immune response (6). DNA vaccines are heat-stable that makes the storage and transport of these vaccine easy (7). Preclinical efficacy of DNA vaccines has been demonstrated in animal models against SARS (8), MERS (9), influenza virus, human immunodeficiency virus, rabies virus, hepatitis B virus, lymphocytic choriomeningitis virus, malarial parasitesand mycoplasma (10, 11).

Studies have reported higher immune response of DNA vaccine by their administration using spring-powered and needle-free devices in animals such as mice, chickens, pigs and non-human primates (NHP) (12–17). The delivery procedure through skin is less painful and it produces higher immunogenicity because of abundance of antigen-presenting cells and dermal lymphatic vessels in skin (18).

Currently, there are twenty-four DNA-based vaccine candidates developed by different pharmaceutical/biotech companies. Sixteen of themare in pre-clinical stage, while eight have progressed toclinical trials (3). ZyCoV-D DNA vaccine candidate developed by Cadila Healthcare Limited has recently completed phase I/II clinical trial (CTRI/2020/07/026352). Here, we report the development and evaluation of the immunogenicity and protective efficacy of ZyCoV-D DNA vaccine candidates in rhesus macaques against SARS-CoV-2 infection.

## RESULTS

### Anti-SARS-CoV-2 IgG response

We evaluated anti-SARS-CoV-2 Immunoglobulin-G (IgG) antibody response against the spike proteinS1 region from the macaque serum samples during the immunization phase (0, 28, 42, 56,70, 84 and 103 days) and SARS-CoV-2 challenge [0, 1, 3, 5, 7, 11 and 15 days post infection (DPI)]. During the immunization phase, animals of group II (1 mg /NFIS) and group IV (2 mg /NFIS) elicited IgG (S1) response starting from day 42 and a gradual rise till day 103 (Fig. 1. A and C). Animals of group III (2 mg /syringe needle) had detectable IgG (S1) levels starting at day 28, but the response was not significanttill day 103. Group I animals (1 mg /syringe needle) and group V (placebo) did not show IgG (S1) response throughout the period of immunization phase (Fig. 1. A and C). After the virus challenge, animals of group IV continued to have higher IgG (S1) titers and whichincreasedfurther post challenge from 7 DPI onwards. Animals of group II and group III had detectable levels of IgG (S1) antibodies from 0 to 7DPI, albeit lower in comparison to group IV. Group I and group V animals had no detectable titers of IgG (S1) till 7 DPI (Fig. 1.B and D). By 15 DPI, animals of all the groups had developed IgG (S1) antibodies as a result of the virus challenge, but titers of group IV wassignificantly higher (1:3200) as compared to other groups (Fig. 1 B and E).

**Fig. 1.**
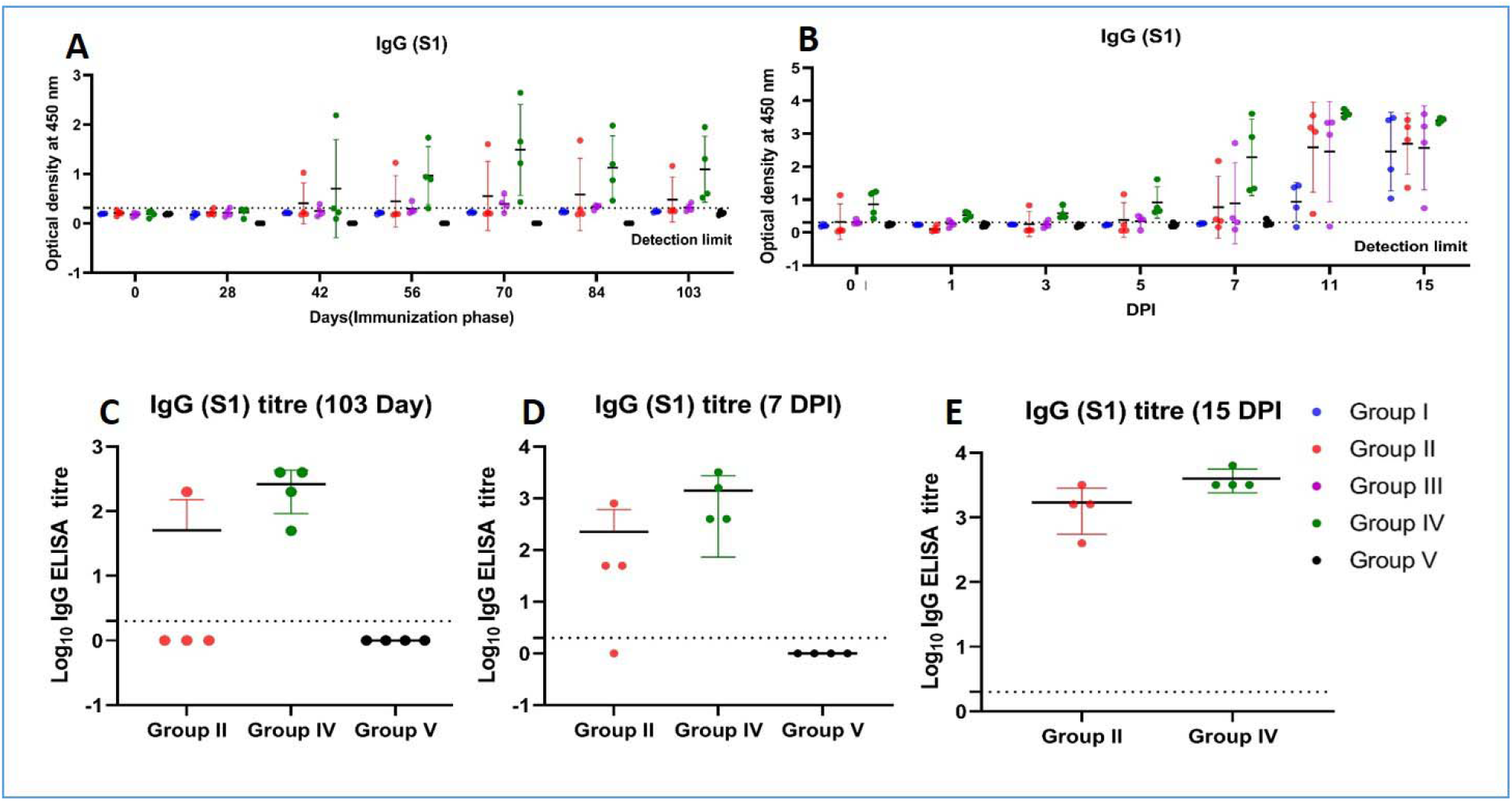
Anti-SARS-CoV-2 spike (S1) gene IgG response in rhesus macaques. (A) IgG response in animals of group I, II, III, IV and V from 0 to 103 days of immunization period (B) IgG response in animals of group I, II, III, IV and V at 0, 1, 3, 5, 7, 11 and 15 DPI (post challenge) (C) Peak IgG titers at day 103 of immunization period of group II, IV and V (D) Peak IgG titers at 7 DPI of group II, IV and V (E) Peak IgG titers at 15 DPI of group II and IV. The statistical significance was assessed using the Kruskal-wallis test followed by the two tailed Mann-Whitney test between two groups; p-values of less than 0.05 were considered to be statistically significant. The dotted line on the figures indicates the limit of detection the assay. Data are presented as mean values +/-standard deviation (SD). Statistical comparison was done by comparing the vaccinated group with the placebo group as control. Group I = blue, group II = orange, group III = purple, group IV = green and group V = black, number of animals = 4 animals in each group.

### Neutralizing antibody response

Neutralizing antibody (NAb) titers of serum samples of animals of all groups were determined during the immunization phase (0, 28, 42, 56,70, 84 and 103 days) and post SARS-CoV-2 challengeat 0, 1, 3, 5, 7, 11 and 15 DPI for group I-IV, while for group V animals it was performed till 7 DPI. Group IV animals had the earliest appearanceof NAb (day 42) which increasedtillday 103. Animals of group II (1/4) and group III (1/4) too developed NAb, but had lower titers as compared to group IV. Animals of group I and group V did not develop NAb during the immunization period till day 103 (Fig. 2. A and C). After the virus challenge, animals of group IV continued to have higher NAb titers as compared to group II and III till 7 DPI. Animals of group I and group V did not develop NAb till 7 DPI (Fig. 2. B and D). By 15 DPI, all the animals of group I–IV developed NAb and animals of group IV showed significantly high titers (1:1452 to 1:3296) (Fig. 2. B and E).

**Fig. 2.**
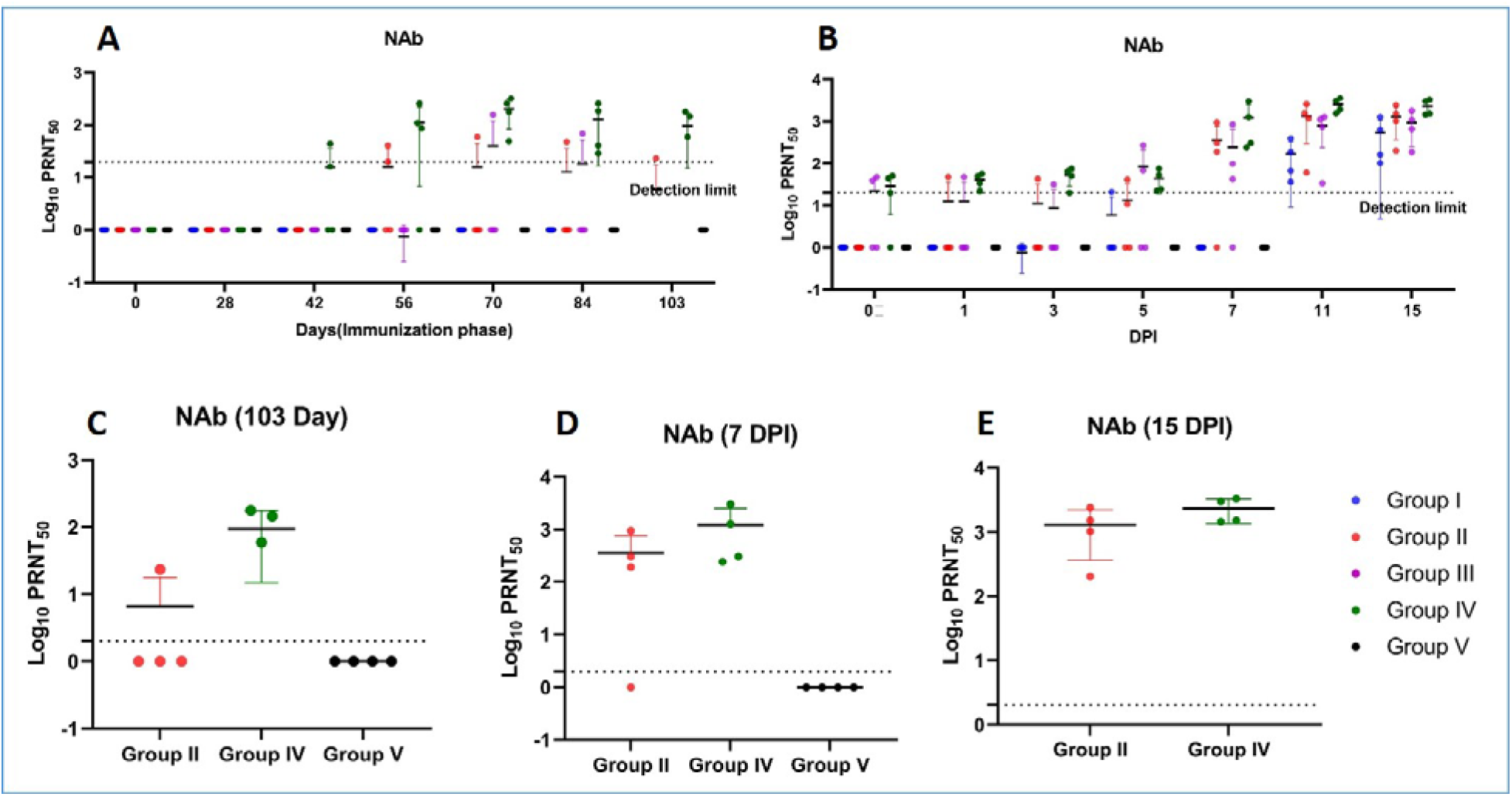
Neutralizing antibody response in rhesus macaques. (A) NAb titers in animals of group I, II, III, IV and V from 0 to 103 days of immunization period (B) NAb titers in animals of group I, II, III, IV and V at 0, 1, 3, 5 7, 11 and 15 DPI (C) Peak NAb titers at day 103 of immunization phase of group II, IV and V (D) Peak NAb titers at 7 DPI of groups II, IV and V (E) Peak NAb titers at 15 DPI of group II and IV. The statistical significance was assessed using the Kruskal-wallis test followed by the two tailed Mann-Whitney test between two groups; p-values of less than 0.05 were considered to be statistically significant. The dotted line on the figures indicates the limit of detection the assay. Data are presented as mean values +/-standard deviation (SD). Statistical comparison was done by comparing the vaccinated group with the placebo group as control. Group I = blue, group II = orange, group III = purple, group IV = green and group V = black, number of animals = 4 animals in each group.

### Viral load in nasal swab, throat swab and bronchoalveolar lavage fluid

Among the vaccinated groups, group IV showed higher rate of reduction in viral load in both NS and TS samples up to 7 DPI. Virus cleared in all animals by 11 DPI except in NS of one animal in group II which also cleared of by 15 DPI (Fig. 3. A and B).

**Fig. 3.**
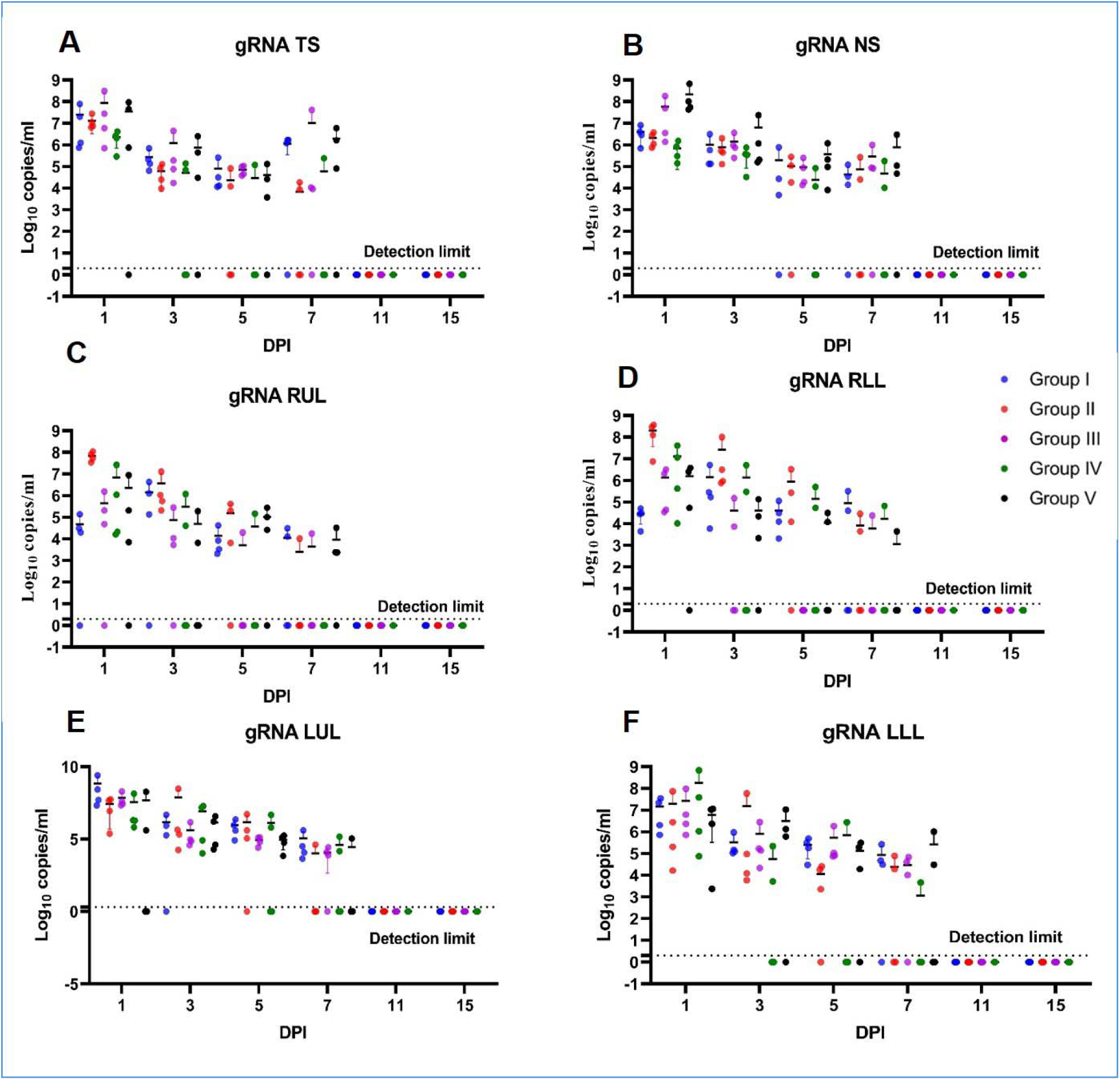
Genomic viral RNA detection in respiratory tract specimens of animals. Genomic viral RNA load in (A) throat swab (TS) (B) nasal swab (NS) (C) right upper lobe (RUL) BAL (D) right lower lobe (RLL) BAL (E) left upper lobe (LUL) BAL (F) left lower lobe (LLL) BAL at 1, 3, 5, 7, 11 and 15 DPI. The statistical significance was assessed using the Kruskal-wallis test followed by the two tailed Mann-Whitney test between two groups; p-values of less than 0.05 were considered to be statistically significant. The dotted line on the figures indicates the limit of detection the assay. Data are presented as mean values +/-standard deviation (SD). Statistical comparison was done by comparing the vaccinated group with the placebo group as control. Group I = blue, group II = orange, group III = purple, group IV = green and group V = black, number of animals = 4 animals in each group.

Group IV animals also showed faster viral clearance from the lung lobes as compared to animals from other groups at 7 DPI (Fig. 3. C, D, E and F). Complete viral clearance was seen in all animals of vaccinated groups by 15 DPI (Fig. 3. A-F).

Subgenomic RNA (sgRNA) was detected in NS of 2/4 of group V at 1 DPI. TS of group V (2/4 animals) and group I (1/4 animals) also had detectable level of sgRNA at 1DPI. All the animals of group I, II, V and 2/4 of the animals from group IV had presence of sgRNA in BAL fluid at 1 DPI. sgRNA was also detected in BAL fluid of group IV and group II in animals till 3 DPI respectively. In group III animals sgRNA was absent in all the specimens from 1 to 7 DPI (Fig. 4 A-F).

**Fig. 4.**
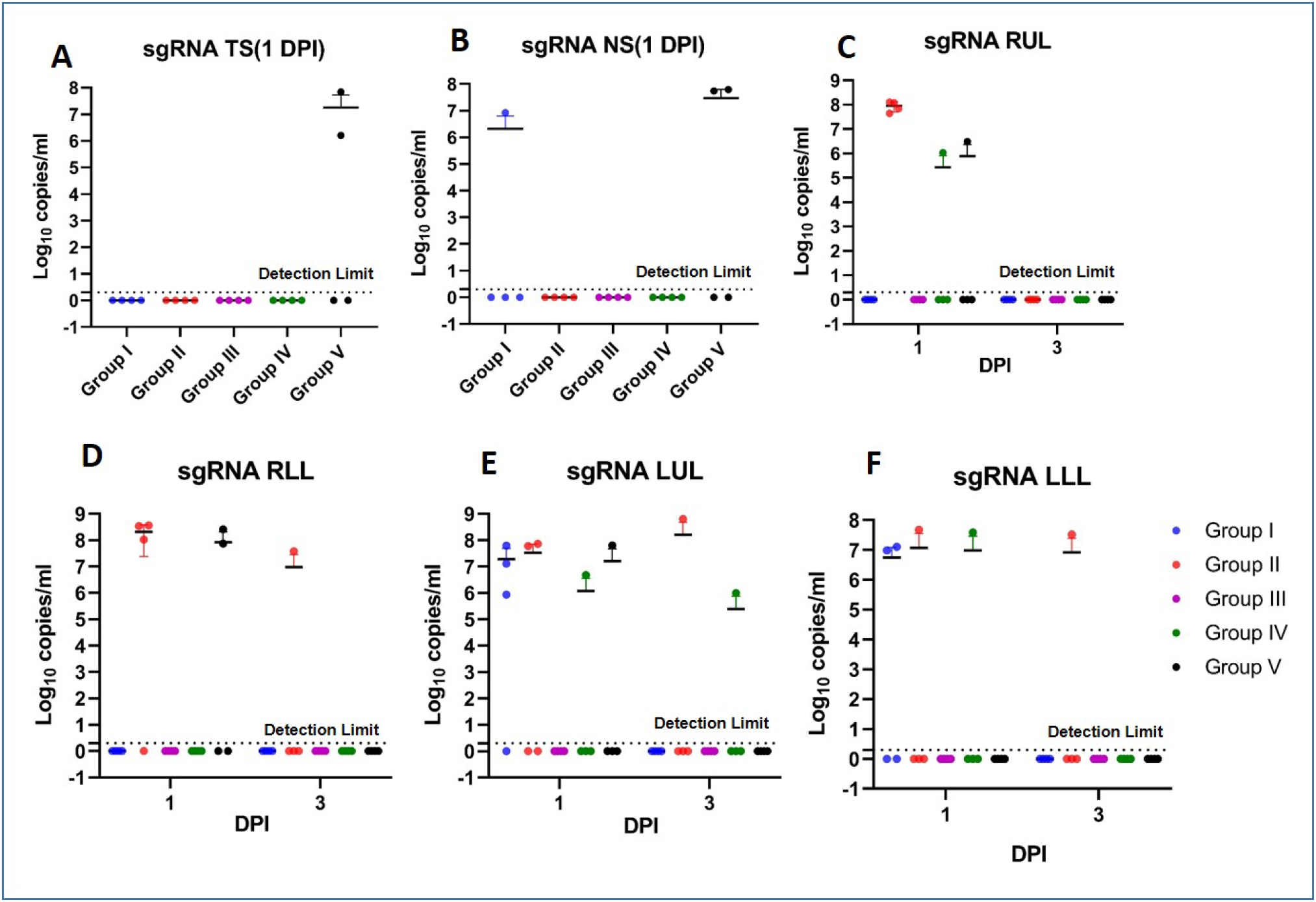
Subgenomic viral RNA detection in respiratory tract specimens of animals. Subgenomic viral RNA load in (A) Throat swab (TS) at 1 DPI (B) Nasal swab (NS) at 1 DPI (C) right upper lobe (RUL) BAL at 1 and 3 DPI (D) right lower lobe (RLL) BAL at 1 and 3 DPI (E) left upper lobe BAL (LUL) at 1 and 3 DPI (F) left lower lobe (LLL) BAL at 1 and 3 DPI. The statistical significance was assessed using the Kruskal-wallis test followed by the two tailed Mann-Whitney test between two groups; p-values of less than 0.05 were considered to be statistically significant. The dotted line on the figures indicates the limit of detection the assay. Data are presented as mean values +/-standard deviation (SD). Statistical comparison was done by comparing the vaccinated group with the placebo group as control. Group I = blue, group II = orange, group III = purple, group IV = green and group V = black, number of animals = 4 animals in each group.

### Viral load in respiratory tract, lungs and extra-pulmonary organs

Animals of group V were euthanized on 7 DPI. Pulmonary and extra-pulmonary organs/tissues were collected. Gross pathology of lung showed broncho-pneumonic patches and consolidation in lungs at necropsy. gRNA was detected in trachea (2/4), nasopharyngeal mucosa (2/4), oropharyngeal mucosa (1/4) and nasal mucosa (2/4) specimens of the sacrificed animals. The animals had detectable gRNA in multiple lobes of lungs, mediastinal lymph nodes (2/4), tonsil (1/4), stomach (1/4) and colon (1/4). Other extra-pulmonary organs (cervical lymph node, heart, liver, kidney, spleen, urinary bladder, small/large intestine, skin and brain) were negative for gRNA.

### Clinico-radiological analysis

Body temperature and body weights of animals in all groups remained within normal range throughout the post challenge period. Hydration status and hair coat condition remained normal in animals of all the groups till 15 DPI. No nasal discharge or lacrimation was observed in any of the animals till 15 DPI. RR was found to increase in 3/4 animals of group I, 2/4 animals of group II, 1/4 animals in group III and 2/4 animals of group IV and 1/ 4 macaque of group V at 7 DPI. SpO_2_ was observed to be lower in 2/4 animals of group V, 3/4 animals of group 2 and 4/4 animals of group IV at 3 DPI.

Fifty percent (2/4) animals of group V developed radiological lesions on chest X-ray (CXR) suggestive of consolidation by 3 DPI which persisted till the 7 DPI (Fig.5 A).

**Fig. 5.**
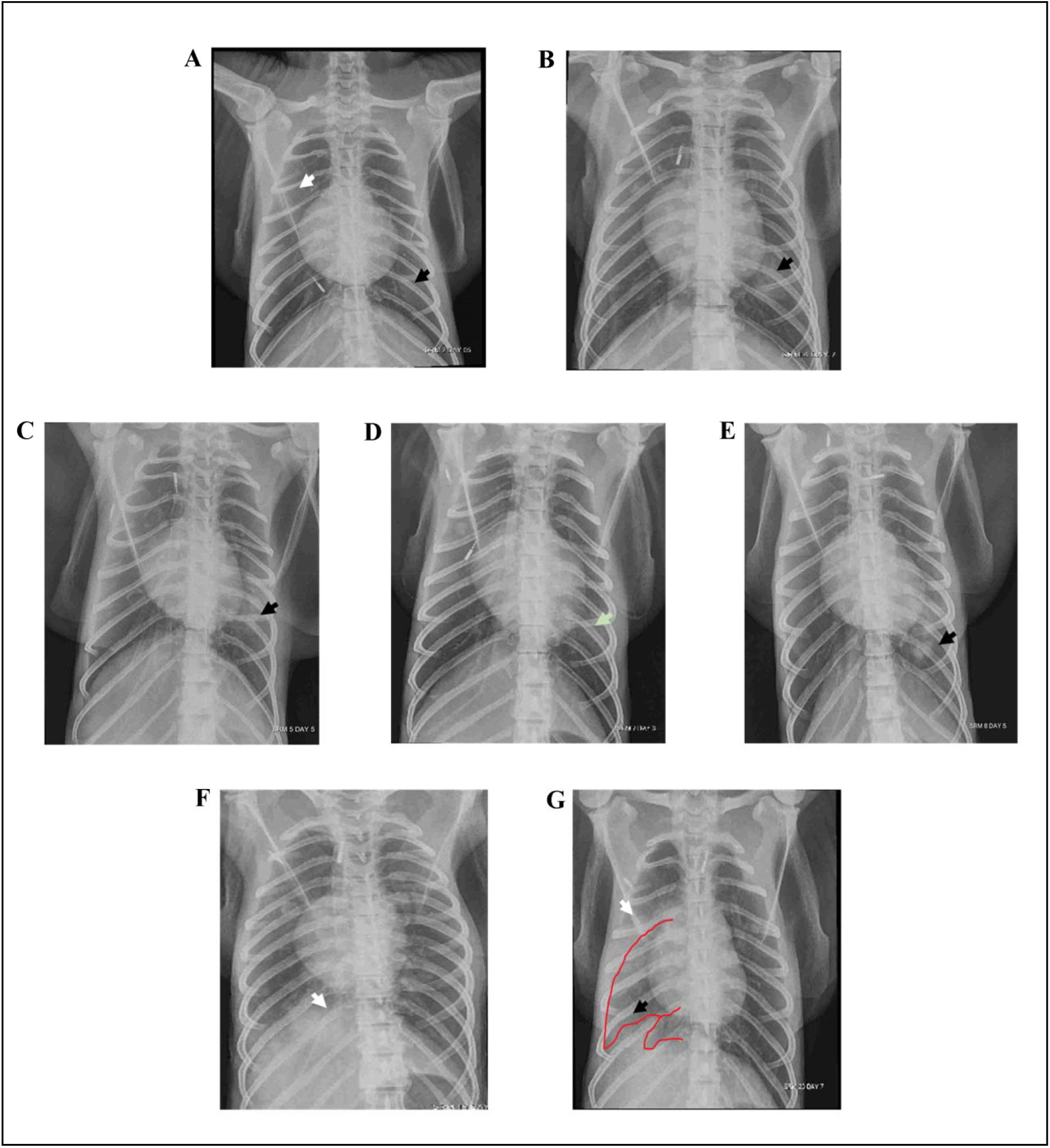
X-rays depicting the radiological lesions in animals of group V, I and IV. (A) Bilateral lung involvement in the form of lobar pneumonia of the right upper and left lower lobes in an animal of group V at 5 DPI (B) Left lower lobe pneumonia in an animal of group V at 7 DPI (C) Lobar pneumonia of the left lower lobe noted on 3 DPI which wa noted to be resolving by 5 DPI in an animal of group I (D) Lobar pneumonia of the Left middle lobe noted on 3 DPI in an animal of group I (E) Lobar pneumonia of the left lower lobe noted on 5 DPI in an animal of group I (F) Lobar consolidation of the right lower lobe noted on 7 DPI in an animal of group IV (G) Lobar consolidation of the right upper and lower lobes, associated with ipsilateral pleural effusion was noted on 7 DPI in an animal of group IV

Radiological lesions suggestive of consolidation appeared on 3 DPI in two and 5 DPI in one animal of group I respectively. In the two animals in which the lesions were seen on 3 DPI, the same were in the form of lobar consolidation of the LLL and LML respectively and resolved by 7 DPI (Fig.5 B).

No radiological lesions have been observed in any of the animalsfrom Group II and III till 15 DPI. Two animals from Group IV displayed radiological features consistent with lobar consolidation/infiltrates (RLL in one animal and RUL/RLL in the other) that appeared on the 7DPI. The lung lesions in the two animals of group IV resolved by 11 DPI / 13 DPI (Fig.5 C). Chest X-ray evaluation in the post challenge period reveals that lesions appeared in group I and group V early in the post challenge period (3 DPI onwards) and persisted till 7 DPI with gradual resolution later. However, in group IV lesions appeared late (7 DPI) and resolved by 11/13 DPI.

### Assessment of lymphocyte subsets

Post-challenge with the virus, the percentages of peripheral NK cells were significantly lower in all vaccinated groups and in the placebo group except for group IV on 1 DPI compared to 0 DPI. NK cell response rose to become significantly higher in vaccinated group III and in group IV at 7 DPI and 15 DPI onwards respectively (Fig 6B). Post challenge, on 1 DPI the percentages of peripheral B cells were significantly higher in group I and group IV compared to 0 DPI. The percentages of B cells were comparable in all vaccinated groups at 7 and 15 DPI compared to 1 DPI. However, in group III, B cells were significantly elevated at 7 DPI Vs. 15 DPI (Fig 6C). The percentages of peripheral CD4 +T cells post-challenge were significantly higher in group I, group II and in placebo group on 1 DPI compared to 0 DPI. In group I, the percentages of CD4+T cells significantly decreased at 15 DPI compared to 1DPI. However, CD4+T cells were significantly elevated in group IV at 15DPI compared to both 7 and 1 DPI (Fig 6D).

**Fig. 6.**
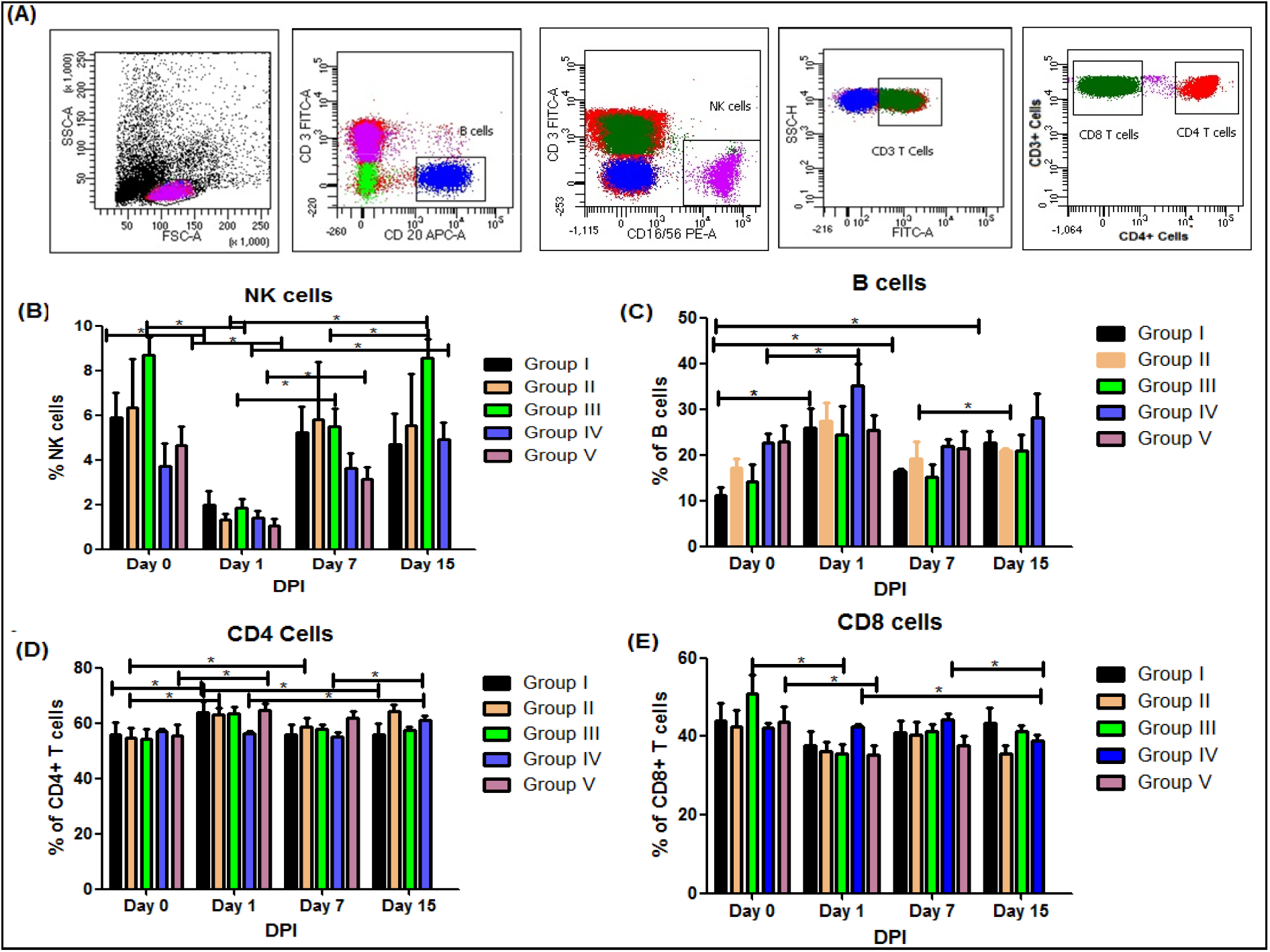
The percentages of lymphocytes in vaccine candidate immunized monkeys: (A) Gating strategy of identification of B cells, NK cells, CD4+ and CD8+ T cells (B) percentages of NK cells (C) percentages of B cells (D) percentages CD4+ T cells (E) percentages of CD8+ T cells at day 0, 7 and 15 DPI. The statistical significance was assessed using the Kruskal-wallis test followed by the two tailed Mann-Whitney test between two groups; p-values of less than were considered to be statistically significant. The dotted line on the figures indicates the limit of detection the assay. Data are presented as mean values +/-standard deviation (SD). Statistical comparison was done by comparing the vaccinated group with the placebo group as control. Group I = black, group II = yellow, group III = green, group IV = blue and group V = pink, number of animals = 4 animals in each group.

The percentages of peripheral CD8+T cells post-challenge were significantly lower in group III and in placebo groups on 1 DPI compared to 0 DPI. However, the percentages of CD8+T cells remained significantly lower in group IV at 15 DPI compared to both 1 and 7 DPI (Fig 6E).

### Cytokine/chemokine profile

IL-8, a key chemokine, responsible for the recruitment of neutrophils and other immune cells at the site of infection was substantially high in vaccinated group IV on 1 and 5 DPI indicative of host immune responses to SARS-CoV-2 infection, which subsided on 7 DPI. A recent study on rhesus macaques post challenge with SARS-CoV-2 demonstrated neutropenia, which could be associated with comparatively low IL-8 level in the control group (19).

IL-5, a Th2 cytokine is primarily associated with eosinophilia. There is compelling experimental evidence that eosinophils have potential antiviral activity (20). Our study demonstrated an elevated level of serum IL-5 in vaccinated group II on 0, 1 and 7 DPI and group IV at 7 DPI which subsided by 11 DPI. The pro-inflammatory cytokine, IL-6 was found transiently increased on 1 DPI in all the study groupsdue to SARS-CoV-2 challenge, which subsided thereafter (Fig. 7).

**Fig. 7.**
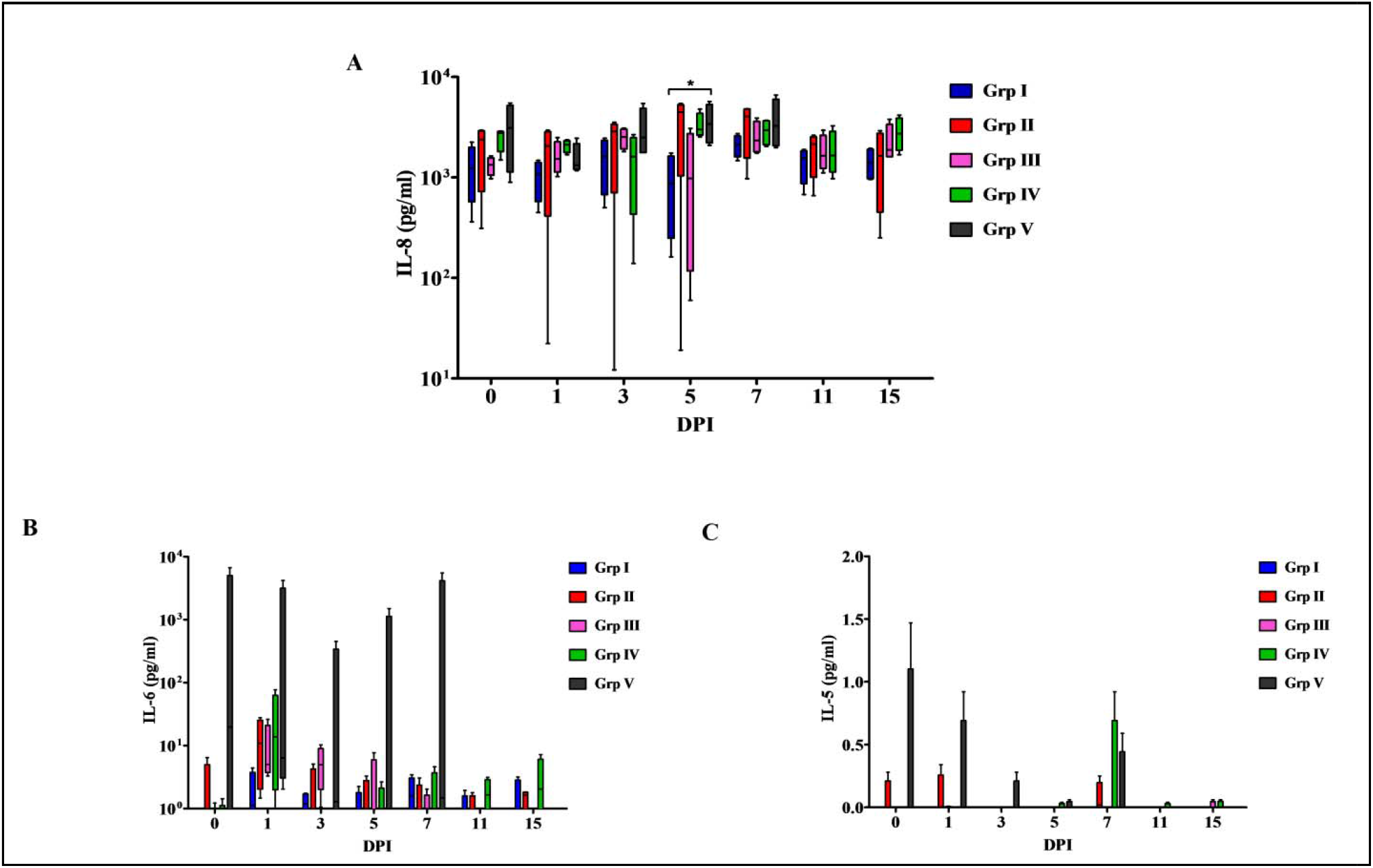
Serum cytokines in rhesus macaques challenged with SARS-CoV-2. (A) IL-8 (B) IL-6 (C) IL-5. The statistical significance was assessed using the Kruskal-wallis test followed by the two tailed Mann-Whitney test between two groups; p-values of less than 0.05 were considered to be statistically significant. The dotted line on the figures indicates the limit of detection the assay. Data are presented as mean values +/-standard deviation (SD). Statistical comparison was done by comparing the vaccinated group with the placebo group as control. Group I = blue, group II = red, group III = pink, group IV = green and group V = black, number of animals = 4 animals in each group.

### Virus isolation

NS, TS and BAL fluid (multiple lobes) of animals from groups I to V; lung lobes and mediastinal lymph node specimens of group V animals were processed for virus isolation (n=265) using Vero CCL-81 cells for two passages (Passage 1 and 2). Cytopathic effect (CPE) was observed in TS, NS, BAL fluid and lung specimens (n=43) of the animals. NS and TS specimens of majority of the animals from all the groups were found to be positive on 1 DPI. NS specimens of animals from vaccinated group I, II and III have also shown CPE on 3 and 5 DPI respectively. Besides this, TS specimen of one animal from group III yielded positive CPE on 7 DPI. However, the virus isolation from animals of group IV was only obtained till 3 DPI (only from TS and NS) suggesting the early clearance of replication competent virus compared to the animals from other groups (Table S1).

## DISCUSSION

SARS CoV-2 pandemic has caused devastating consequences to the public health and economy globally. Development of multiple therapeutic or prophylactic countermeasures is the need of the hour. Many vaccine candidates have progressed to the clinical trials and preclinical research in animal models (21–24). This includes whole virion inactivated, mRNA, DNA, vectored vaccines etc. Among the different vaccine platforms, DNA vaccines appear promising with advantages like stability, efficient large scale low costproduction and ability to induce enhanced humoral and cell mediated immune response (25). Majority of SARS-CoV-2 vaccine studies describe the protection conferred by the vaccination before waning of the acute phase immune response (21, 22, 24). The long-term immunological memory induced by these vaccine candidates yet needs to be explored as studied in case of few candidates (26, 27). This is important as SARS-CoV-2 can cause asymptomatic or mild disease in humans without inducing significant antibody response. In the current study, we evaluated the immunogenicity of ZyCoV-D DNA vaccine candidates in rhesus macaques and the protective efficacy against SARS-CoV-2 infection post 14 weeks of first immunization dose.

ZyCoV-D vaccine candidate (2 mg by NFIS) showed good immunogenicity as evident by the anti-SARS-CoV-2 S1 IgG and NAb titers. The vaccine candidate given as 1mg (NFIS) and 2 mg (syringe needle) could elicit the immune response in only few animals indicating the importance of the dose and vaccine delivery system in evoking the immune response. A dose dependent increase in the IgG and T cell response has been demonstrated with DNA vaccine candidate, GX-19 in NHP (26), which could be the reason for the reduced immunogenicity conferred by the 1mg dose NFIS group. Studies have reported the efficiency of *in vivo* delivery affects in the induction of immune response by a DNA vaccine (26, 28, 29). The better response of 2mg (NFIS) could also be due to the difference in vaccine delivery. Needle-free injectors for DNA vaccine delivery have been found to augment immune response, possibly by enhancing the dispersion of injecting substance or increasing local inflammation (29). Multiple approaches have been tried by various groups for vaccine delivery in case of SARS-CoV-2 DNA vaccines like intramuscular, intradermal, oral and electroporation techniques (23, 26, 27, 30). Use of electroporation method for GX-19 vaccine candidate delivery enhanced the antibody and T cell response in NHP’s (26).

A preclinical study of multiple SARS-CoV-2 DNA vaccine candidates expressing six variants of S protein has reported effective protection of rhesus macaques with significant NAb titre (23). A durable humoral immune response could be observed till 14 weeks with the ZyCoV-D vaccine (2 mg NFIS) which was enhanced in the post challenge and persisted till 15 DPI as compared to other vaccinated groups. The 2 mg dose injected by NFIS also showed detectable level of NAb titer from 6 weeks with an incremental response throughout the immunization phase.

INO-4800, GX-19 and a full-length S protein DNA vaccine candidate showed NAb response by 4, 5.5 and 5 weeks, which persisted till 12, 8 and 6 weeks respectively following which animals were challenged with the virus (23, 26, 27). Other SARS-CoV-2 vaccines which progressed to clinical trials like inactivated vaccines (PicoVacc, BBV152), mRNA vaccines (mRNA-1273, BNT162b2, ARCoV), viral vector vaccines (Ad26.COV2.S and ChAdOx1 nCoV-19), protein sub unit vaccines (NVX-CoV2373, S-Trimer, RBD, Sad23L-nCoV-S/Ad49L-nCoV-S and S1-Fc) showed an early NAb in preclinical studies (31).

Although the complete viral clearance was observed in the ZyCoV-D (2mg by NFIS) vaccinated animals by 14 days, reduction in viral loads were observed in the TS and NS specimens by 7 days post challenge compared to the unvaccinated macaques demonstrating the protective efficacy. The other DNA vaccine candidate’s studies like INO-4800 and GX-19 in macaques were terminated by 7 and 4 DPI respectively. Reduced viral loads in the NS and TS were observed in GX-19 vaccinated macaques four days post challenge (26). Similarly, significantly lower peak sgRNA loads and viral RNA loads in NS and BAL were observed in INO-4800 vaccinated macaques by 7 DPI.

A limitation of our study is that the control animals were followed only till 7 days. Viral clearance was observed from ZyCoV-D vaccinated macaques starting from week 1 and complete clearance was observed by second weeks (31).

There was significant increase in the percent lymphocytes and IL-8 in group IV suggestive of the vaccine induced host immune response/lymphoproliferation to infection. The present study indicated the transient decrease in the % CD8+ T cell population on 1 DPI and 3 DPI. By now, it is well established that lymphopenia due to SARS-CoV-2 infection is biased towards CD8+ T cells. A recent study on rhesus macaque post challenge with SARS-CoV-2 demonstrated neutropenia, which could be associated with comparatively low IL-8 level in the control group (19). Our study demonstrated an elevated level of serum IL-5, a Th2 cytokine is primarily associated with eosinophilia in group II and IV, which subsided by 11 DPI. There is compelling experimental evidence that eosinophils have potential antiviral activity. Other DNA vaccine candidates like GX-19 produced a Th1 cell response with TNF-α, IFN-γ & IL-2 in vaccinated macaques whereas INO-4800 induced only IFN-γ (26, 27).

In conclusion, the ZyCoV-D vaccine candidate, 2mg by NFIS elicited significant SARS-CoV-2 specific IgG, NAbtiters and lesser viral loads in animals post challenge demonstrating its protective efficacy.

## MATERIALS AND METHODS

### Ethical review

The study was approved by the Institutional Project Review Committee, and Institutional Biosafety Committee, ICMR-National Institute of Virology (NIV), Pune. The study was recommended by theInstitutional Animal Ethics Committee of Zydus Research Centre (Registration 77/PO/RcBi/SL/99/CPCSEA) and also by Institutional Animal Ethics Committee of ICMR-NIV (Registration 43/GO/ReBi/SL/99/CPCSEA) and further approved by Committee for the Purpose of Control and Supervision of Experiments on Animals (CPCSEA), New Delhi (Letter no. V-11011(13)/15/2020-CPCSEA-DADF dated 5th October 2020). Similarly, Cadila Healthcare Limited also got the approval of the study protocol (VTC/R&D/nCoV/NHP-01) from CPCSEA. The research was conducted in compliance with the guidelines laid down by CPCSEA.

### Generation of vaccine

The DNA vaccine candidate against SARS-CoV-2 is comprised of a DNA plasmid Vector pVAX1 carrying spike-S protein of SARS-CoV-2. A chemically synthesized Spike region was inserted into pVAX-1 plasmid vector. Plasmid DNA carrying Spike gene was chemically transformed into E coli cells to select the desired clone. The E coli carrying the plasmid DNA was further propagated for large scale production. ZyCoV-D vaccine contains purified concentrated DNA formulated in Phosphate buffered saline having no preservative (32).

### Study design

Twenty healthy, rhesus macaques (*Macaca mulatta*) aged 7-12 years from Primate Research Facility of Cadila Healthcare Ltd, Zydus Research Centre, Ahmedabad were selected for the study which were further divided into 5 groups of 4 animals (2 M, 2 F) each viz., vaccinated group (I, II, III, IV) and placebo group (V).

The animals were immunized at Zydus Research Centre Ahmedabad and further virus challenge experiments were conducted at ICMR-NIV, Pune. The animals were immunized with vaccine candidates,1 mg syringe and needle (intradermal) (group I), 1mg by NFIS (group II), 2mg syringe and needle (intradermal) (group III) and 2mg by NFIS (group IV). The animals were immunized on day 0, 28 and on day 56. Intradermal injections were given in the forearms. The dose volume for each animal did not exceed 0.1ml/site. Blood samples were collected on day 0, 28, 42, 56, 70 84 and 103 for serological analysis. A placebo group of 4 animals (2 male and 2 female) was also included in the study (Fig. 8). Blood samples for humoral response (IgG (S1) and NAb), cytokine/chemokine profile and cell phenotyping were collected on day 90 and 103 before the initiation of virus challenge studies.

**Fig. 8.**
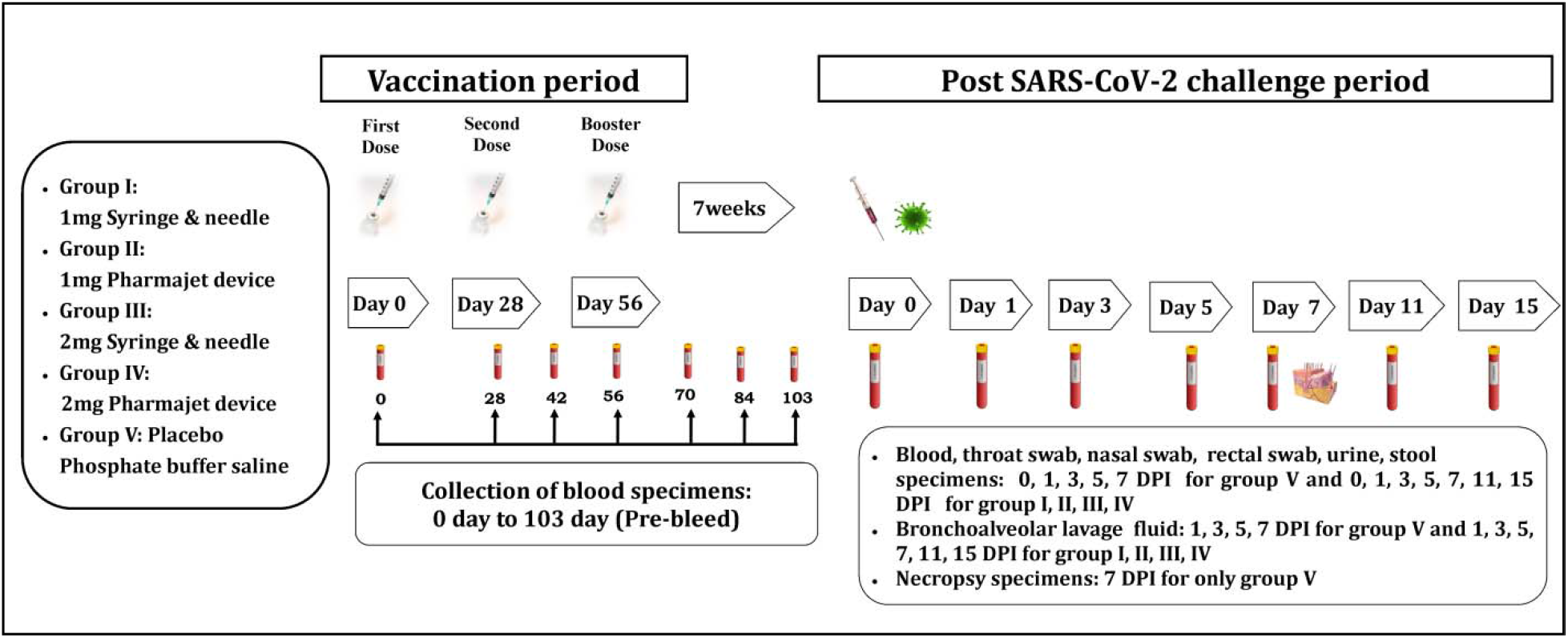
Diagrammatic representation of the study design. Note: Group V animals were euthanized on 7 DPI. Hence in data analysis figures of variables evaluated are shown for group I – V till 7 DPI and group I – IV till 15 DPI

Animals were observed daily for clinical signs and tested for SARS-CoV-2 by real time RT-PCR, haematology (complete blood count, haemoglobin, PCV, DLC) and serum biochemistry prior to initiation of challenge phase. Serum biochemical parameters monitored were totalbilirubin, ALT, AST, ALP, g-glutamyl transpeptidase, total protein, albumin, triglycerides, total cholesterol, creatinine, uric acid, blood urea, creatine kinase and lactate dehydrogenase. The animals of all groups were challenged with 10^6.5^ TCID_50_/ml of SARS-CoV-2 (total volume = 1.5ml) through the intranasal route (0.25ml/ each nostril) and intratracheal route (1ml) at 15 week.

Animals were observed for body temperature, body weight, PR, RR, oxygen saturation at room air (SpO_2_) on 1, 3, 5, 7, 11 and 15 DPI. Clinical signs such as cough, nasal discharge, lacrimation, respiratory distress, lethargy, diarrhoea, ruffled fur coat and rash twice daily from 0 to 15 DPI (Table S2). Blood, NS, TS, RS, urine and stool specimens were collected from each animal on 0, 1, 3, 5, 7, 11 and 15 DPI. BAL fluid aspiration from four lobes of lungs [right upper lobe (RUL), right lower lobe (RLL), left upper lobe (LUL) and left lower lobe (LLL)] was performed on 1, 3, 5, 7, 11 and 15 DPI using a flexible paediatric bronchoscope mm) (Pentax Medical India Private Limited under general anaesthesia. Chest X-ray was done on 0, 1, 3, 5, 7, 11 and 15 DPIto monitor radiological evidence of lung pathology.

### Detection of SARS-CoV-2 genomic RNA and subgenomic RNA

RNA was extracted from NS, TS, RS, BAL, urine, stool and homogenized necropsy specimens using MagMAX ™ Viral/Pathogen Nucleic Acid Isolation Kit (Thermo Fisher Scientific, USA) following the manufacturer’s instructions. Real-time RT-PCR for SARS-CoV-2 genomic RNA (gRNA) and subgenomic RNA (sgRNA) targeting the E-gene was performed on extracted RNA samples as described earlier (33, 34).

### Detection of anti-SARS-CoV-2 IgG antibodies using Spike-protein based-ELISA

Ninety-six well microtiter plates were coated with 100 µL/well of diluted S1 protein (1µg/ml with Phosphate buffer saline) incubated for 16-18 hours at 2-8°C. Plates were washed 3 times with phosphate-buffered saline with 0.05% Tween 20 (PBST) to remove any unbound antigen and blocked with blocking buffer (2.5 g of skimmed milk in 1X PBS) for one hour at 37°C. The wells were washed 3 times with PBST and were incubated at 37°C for one hour with 100μL of diluted macaque serum samples (1: 50). Negative control was added to each plate. After 5 washes with PBST, anti-monkey IgG HRP-antibodies 1:5000 (Thermoscientific, USA) were added and incubated for 1 hour at 37°C. Following 5 washes with PBST, 100 μL of 3’,3’5,5’-tetramethylbenzidine (TMB) substrate was added to each well and incubated for 15-20 minutes in dark. The reaction was terminated by adding 100μL of the stop solution to each well and absorbance was measured at 450 nm after 10 minutes. Serum IgG titers were determined by testing serial 10-fold dilutions of each sample, starting from 1:50 dilution.

### Detection of neutralizing antibodies using PRNT

The plaque reduction neutralizing antibody test (PRNT) was performed using the protocol described earlier (35). Briefly, four-fold serial dilutions of serum samples were mixed with an equal volume of virus suspension containing 50-60 plaque-forming units (PFU/0.1ml). Post an hour of incubation at 37°C, the virus–serum mixtures (0.1ml) were added onto the Vero CCL-81 cell monolayers and incubated at 37°C with 5% CO_2_for 1 hour. Subsequently, the suspension was aspirated and the cell monolayer was gently overlaid with an overlay medium containing 2% carboxymethylcellulose (CMC) and 2X Minimum essential medium (MEM) containing 2% Foetal bovine serum (FBS) (1:1) and the plates were further incubated at 37°C with 5% CO_2_ for 5days. The plates were stained with 1% amido black and plaque numbers were counted. PRNT_50_ titer was calculated using a log probit regression analysis by SPSS (SPSS Inc., Chicago, IL).

### Cytokine/chemokine and growth factor level

The levels of IL-2, IL-5, IL-6, IL-8, IFN-γ and TNF-α in the serum samples were analysed using BD cytometric bead array flex sets, as per manufacturer’s instructions and as previously described (36). Briefly, 50µl serum sample was incubated with capture beads coupled with antibodies. The immune complexes were incubated with detection antibody conjugated with phycoerythrin. The standards provided by the manufacturer were used to prepare the standard curve for each cytokine. Cytokine levels were measured on a BD FACSCaliburTM flow cytometry (BD Biosciences) using BD CellQuestTM Pro software. The data was analysed using flow Cytometric analysis program (FCAP) ArrayTM software.

### Immunophenotyping of PBMCs

For surface phenotyping, freshly isolated PBMCs from immunized/ SARS-CoV-2 challenged NHPs adjusted to 1 × 10^6^ cells per test were incubated with anti NHP T/B/NK cocktail and anti CD4 PECF594 monoclonal antibodies for 30 min at 4^°^C. This was followed by washing of the cells with washing buffer at 1500 rpm for 10 minutes. The cells were re suspended in 350 μl 1 % paraformaldehyde solution. Stained PBMCs were assessed for the expression patterns of the above immune cell markers by flow cytometry (FACS Aria II, BD Biosciences, USA). For each cell type, 100,000 events were acquired with appropriate isotype control and data were analyzed using FACS Diva software. The gating strategy is depicted in Fig 6A.

### Virus isolation from clinical and necropsy specimens

Lung tissues were homogenized in sterile Minimum Essential Medium (MEM; Gibco, USA) using a homogenizer (GenoGrinder 2000; BT&C Inc., Lebanon, NJ, USA). Further, tissue homogenates were centrifuged at 1984 g for 10 min and supernatant was aliquoted in cryovials. Vero CCL-81 cells were grown to confluent monolayer in 24-well plate maintained in MEM supplemented with 10 %FBS (HiMedia, Mumbai), penicillin (100 U/ml) and streptomycin (100 mg/ml). After decanting the growth medium, 100μl volume of homogenized supernatant and clinical specimen was inoculated on to 24-well cell culture monolayer of Vero CCL-81. The cells were incubated for one hour at 37°C to allow virus adsorption, with rocking every 10 min for uniform inoculum distribution. After the incubation, the inoculum was removed and the cells were washed with 1X PBS. The MEM supplemented with two percent FBS was added to each well. The cultures were incubated further in incubator at 37°C with 5% CO2 and observed daily for cytopathic effects (CPEs) under an inverted microscope (Nikon, Eclipse Ti, Japan). Cultures that showed CPE were centrifuged at 4815 × g for 10 min at 4°C; the supernatants were processed immediately or stored at −80°C (37). The presence of the virus in the culture supernatant was confirmed by SARS-CoV-2 specific real-time RT-PCR.

### Histopathological examination and Immunohistochemistry

Gross examination of lungs was done to assess the degree and extent of consolidation and congestion as per the recommendation of the Working Group of the Society of Toxicologic Pathology’s Scientific and Regulatory Policy Committee (38). Histopathological examination of various lung lobes was carried out to assess the extent of interstial pneumonia, alveolar damage, haemorrhages, inflammatory cell infiltration, hyaline membrane formation and accumulation of eosinophilic oedematous exudate using the procedure as described earlier (39, 40). Immunohistochemistry (IHC) was performed on the lung tissue to detect the presence of SARS-CoV-2 antigen on type-II pneumocytes and alveolar macrophages using the techniques described elsewhere (data not shown here) (39).

### Data analysis

Clinical, virological, serological and immunological data were analysed using GraphPad Prism software version 8.4.3 (GraphPad, San Diego, California). The data for the vaccinated groups were initially compared with the control group using the non-parametric Kruskal-Wallis test. If the Kruskal-Wallis test was found to be significant,a group-wise comparison between the control and the other vaccinated groups was performed to assess significance using a two-tailed Mann-Whitney test. The p-values less than 0.05 were considered significant and are marked on the figures. Non-significant values are not depicted in the figures. The detection limits are marked with the dotted lines on the respective figures. The log10 plots below the detection limits are depicted with a value of 1 for the illustration purpose.

## SUPPLEMENTARY MATERIALS

Table S1. Details of virus isolation using Vero CCL-81 (passage 1 and 2) from clinical and necropsy specimens of macaques

Table S2. Day-wise animal observation sheet

## Acknowledgments

Authors gratefully acknowledge the encouragement and support extended by Prof. (Dr.) Balram Bhargava, Secretary to the Department of Health Research, Ministry of Health & Family Welfare, Government of India & Director-General, ICMR, New delhi and Dr. Samiran Panda, Scientist ‘G’ & Head, Epidemiology and Communicable Diseases, ICMR, New delhi. We also acknowledge the support received from and laboratory team which includes Deepak Mali, Savita Patil, Triparna Majumdar, Rajen Lakra, Hitesh Dighe, Ratan More, Kaumudi Kalele, Yash Joshi, Pranita Gawande, Ashwini Waghmare, Annasaheb Suryawanshi, Manoj Kadam, Ganesh Chopade, Sanjay Thorat, Madhav Acharya, Poonam Bodke, Manisha Dudhmal, Tejashree Kore and engineering staff Mayur Mohite, Vishal Gaikwad, Nandkumar Sharma, Animal Facility team members SiddharamFulari and Diagnostic virology group members Rashmi Gunjikar, Darpan Phagiwala, Chetan Patil and Deepika Chowdhary of ICMR-National Institute of Virology, Pune, India.

We sincerely acknowledge Major General Arindam Chatterjee, Commandant, AICTS, Pune, Major General S Hasnain, Commandant, Command Hospital (Southern Command) Pune, India for extending generous support for the study. We would also thank Havaldar Mangesh Aamre, AICTS, Pune for his excellent support in bronchoscopy.

## Author’s Contribution

Conceptualization: PDY, SK, PA, NG, SG,KM

Design and production of vaccine candiadte: MJ, KM, SG, AD, HC, CR, HPR, SP, NS

Planning and execution of the animal experiments: PDY, SK, KA, SLM, DRP, BM, AK.

Laboratory work planning and data analysis: PDY, ASA, GS, AT, DYP, HK, DN, GD, RS.

Execution of the laboratory experiments: GD, HK, RJ, PS, SB, AK, HD, RM, DS

Writing – original draft: PDY, SK, DYP, ASA, SLM

Writing – review andediting: PDY, SK, DYP, ASA, PA, NG

## Data and materials availability

All the data other than those presented in the article are provided in the form of supplementary files.

