## Supplementary material for "Assessment of immunogenicity and protective efficacy of ZyCoV-D DNA vaccine candidates in Rhesus macaques against SARS-CoV-2 infection": Table S1 and Table S2

**Table S1. Details of virus isolation using Vero CCL-81 (passage 1 & 2) from clinical and necropsy specimens of macaques.**

| **Specimens** | **Day** | **Group V** | **Group I** | **Group II** | **Group III** | **Group IV** |
| --- | --- | --- | --- | --- | --- | --- |
| **Throat swab** | Day 1 | **3/4** | **3/4** | **4/4** | **4/4** | **3/4** |
|  | Day 3 | 0/2 | 0/4 | 0/2 | 0/4 | **1/3** |
|  | Day 5 | 0/4 | 0/1 | 0/1 | 0/2 | 0/1 |
|  | Day 7 | 0/2 | 0/3 | 0/0 | **1/2** | 0/1 |
| **Nasal swab** | Day 1 | **4/4** | **3/4** | **3/4** | **4/4** | **1/4** |
|  | Day 3 | 0/4 | **1/4** | **1/4** | 0/4 | 0/3 |
|  | Day 5 | 0/2 | 0/2 | 0/2 | **1/2** | 0/1 |
|  | Day 7 | 0/2 | 0/2 | 0/1 | 0/3 | 0/1 |
| **RUL BAL** | Day 1 | 0/1 | 0/2 | 0/4 | 0/2 | 0/2 |
|  | Day 3 | 0/0 | 0/3 | 0/4 | 0/1 | 0/2 |
|  | Day 5 | 0/1 | 0/0 | 0/3 | 0/0 | 0/1 |
|  | Day 7 | 0/0 | 0/0 | 0/0 | 0/0 | 0/0 |
| **RLL BAL** | Day 1 | 0/1 | 0/1 | 0/3 | 0/2 | 0/2 |
|  | Day 3 | 0/1 | 0/3 | 0/4 | 0/1 | 0/1 |
|  | Day 5 | 0/1 | 0/1 | 0/2 | 0/0 | 0/1 |
|  | Day 7 | 0/0 | 0/1 | 0/0 | 0/0 | 0/0 |
| **LUL BAL** | Day 1 | 0/3 | **1/4** | 0/4 | **1/4** | 0/4 |
|  | Day 3 | 0/2 | 0/3 | 0/3 | 0/3 | 0/3 |
|  | Day 5 | 0/2 | 0/3 | 0/3 | 0/3 | 0/2 |
|  | Day 7 | 0/1 | 0/1 | 0/1 | 0/0 | 0/1 |
| **LLL BAL** | Day 1 | 0/3 | **3/4** | 0/4 | 0/4 | 0/4 |
|  | Day 3 | 0/3 | 0/4 | 0/2 | 0/3 | 0/2 |
|  | Day 5 | 0/4 | 0/3 | 0/0 | 0/4 | 0/1 |
|  | Day 7 | 0/1 | 0/1 | 0/1 | 0/0 | 0/0 |
| **Lung Lobes** | Day 7 | **1/6** | NA | NA | NA | NA |
| **Mediastinal lymph node** | Day 7 | 0/1 | NA | NA | NA | NA |
| **Total positive** | | **8/55** | **11/58** | **8/56** | **11/52** | **5/44** |
|  |  | **43/265** | | | | |

*(RUL= right upper lobe BAL, RLL= right lower lobe BAL, LUL= left upper lobe BAL, LLL= left lower lobe BAL, NA = Not available)

**Table S2: Day-wise animal observation sheet**

| **Group** | **NIV ID** | **Day 0** | **Day 1** | **Day 2** | **Day 3** | **Day 4** | **Day 5** | **Day 6** | **Day 7** | **Day 8** | **Day 9** | **Day 10** | **Day 11** | **Day 12** | **Day 13** | **Day 14** | **Day 15** |
| --- | --- | --- | --- | --- | --- | --- | --- | --- | --- | --- | --- | --- | --- | --- | --- | --- | --- |
| **Group V** | SRM 1 | 0.5 | 0 | 0.5 | 0 | 0.5 | 0 | 0 | 0 | NA | NA | NA | NA | NA | NA | NA | NA |
|  | SRM 2 | 3 | 2.5 | 1.5 | 0 | 0 | 0.5 | 0 | 1 | NA | NA | NA | NA | NA | NA | NA | NA |
|  | SRM 3 | 3 | 2.5 | 0.5 | 4 | 3.5 | 1 | 4 | 0 | NA | NA | NA | NA | NA | NA | NA | NA |
|  | SRM 4 | 2.5 | 3 | 0.5 | 1.5 | 1 | 1 | 7.5 | 0 | NA | NA | NA | NA | NA | NA | NA | NA |
| **Group I** | SRM 5 | 0 | 0 | 1.5 | 0 | 0 | 1 | 0.5 | 0 | 0 | 0.5 | 0 | 0 | 1 | 3 | 0 | 0 |
|  | SRM 6 | 1 | 0.5 | 0.5 | 0 | 0 | 0 | 0 | 0.5 | 0 | 0 | 0 | 4 | 5 | 2 | 1 | 0 |
|  | SRM 7 | 2.5 | 4 | 1 | 0 | 0 | 1 | 0.5 | 1 | 0 | 4 | 1 | 4 | 5 | 3 | 0 | 0 |
|  | SRM 8 | 3 | 2.5 | 0.5 | 0.5 | 0.5 | 2 | 0 | 1.5 | 0 | 1 | 1 | 3 | 2 | 0 | 0 | 0 |
| **Group III** | SRM 9 | 0 | 0 | 0.5 | 3 | 2 | 0.5 | 0.5 | 0.5 | 0 | 0.5 | 1 | 1 | 0 | 0 | 0 | 0 |
|  | SRM 10 | 0 | 0 | 0.5 | 1.5 | 1.5 | 4.5 | 2 | 1 | 3 | 1 | 1 | 0 | 0 | 0 | 0 | 0 |
|  | SRM 11 | 0 | 0 | 0.5 | 0.5 | 2 | 3.5 | 1.5 | 1 | 2 | 0 | 1 | 0 | 0 | 0 | 0 | 0 |
|  | SRM 12 | 0 | 0 | 0.5 | 0 | 1 | 3.5 | 1 | 1 | 1 | 3 | 1 | 0 | 0 | 0 | 0 | 0 |
| **Group II** | SRM 13 | 0 | 0 | 2 | 0 | 0 | 0 | 1 | 1 | 1 | 1 | 0 | 0 | 0 | 0 | 0 | 0 |
|  | SRM 14 | 0 | 0 | 3 | 1 | 2 | 0 | 0 | 0 | 0 | 1 | 0 | 0 | 0 | 0 | 0 | 0 |
|  | SRM 15 | 0 | 7 | 5 | 2 | 3 | 2 | 0 | 1 | 1 | 1 | 3 | 1 | 0 | 0 | 0 | 0 |
|  | SRM 16 | 0 | 5 | 1 | 2 | 4 | 2 | 1 | 1 | 1 | 1 | 2 | 2 | 3 | 0 | 0 | 0 |
| **Group IV** | SRM 17 | 0 | 2 | 1 | 2 | 3 | 2 | 4 | 4 | 4 | 2 | 1 | 0 | 0 | 0 | 0 | 0 |
|  | SRM 18 | 0 | 1 | 1 | 2 | 2 | 0 | 2 | 2 | 1 | 1 | 1 | 1 | 1 | 1 | 0 | 0 |
|  | SRM 19 | 0 | 4 | 8 | 7 | 7 | 7 | 7 | 7 | 4 | 3 | 2 | 5 | 4 | 0 | 0 | 0 |
|  | SRM 20 | 0 | 2 | 6 | 6 | 5 | 2 | 6 | 7 | 1 | 1 | 3 | 1 | 3 | 1 | 1 | 0 |

*NA= Not applicable

Two observations were made in a day, mean of two observation can give value of 0.5 for example SRM-1 observed to be restless and uncomfortable in morning and was normal in the evening score for the day (1+0)/2 =0.5
